# Membrane potential dynamics unveil the promise of Bioelectrical Antimicrobial Susceptibility Testing (BeAST) for anti-fungal screening

**DOI:** 10.1101/2023.10.05.561038

**Authors:** Tailise Carolina de Souza-Guerreiro, Letícia Huan Bacellar, Thyerre Santana da Costa, Ljubica Tasic, Munehiro Asally

## Abstract

Membrane potential is a useful marker for antimicrobial susceptibility testing (AST) due to the fundamental characteristic of vital cells. However, the difficulties associated with measuring the membrane potential in microbes restrict its broad application. In this study, we present Bioelectrical AST (BeAST) using the model fungus *Saccharomyces cerevisiae*. Using fluorescent indicators (DiBAC4(3), ThT and TMRM), we measured plasma and mitochondrial membrane-potential dynamics upon electric stimulation. We find that a 2.5-second electric stimulation induces hyperpolarisation of plasma membrane lasting 20 minutes in vital *S. cerevisiae*, but depolarisation in inhibited cells. The numerical simulation of FitzHugh-Nagumo model successfully recapitulates vitality-dependent dynamics. The model also suggests that the magnitude of plasma-membrane potential dynamics (PMD) correlates with the degree of inhibition. To test this prediction and to examine if BeAST can be used for assessing novel anti-fungal compounds, we treat cells with biogenic silver nanoparticles (bioAgNPs) synthesised using orange fruit flavonoids and Fusarium oxysporum. Comparing BeAST with optical density assay with various stressors, we show that PMD correlates with the severityof growth inhibitions. These results suggest that BeAST holds promise for screening anti-fungal compounds, offering a valuable approach to tackling antimicrobial resistance.

## Introduction

Living cells maintain their internal ionic condition to be distinct from their external space. For example, potassium is highly accumulated inside cells in almost all species (Danchin and Nikel, 2019). The ionic difference across the membrane gives rise to an electrical potential gradient, known as membrane potential (Nicholls; and Ferguson, 2013). The membrane potential is a component of ion motive force (IMF) that provides a source of free energy for cells to do various chemical and mechanical work, such as membrane transport and ATP synthesis (Benarroch and Asally, 2020; Galera-Laporta et al., 2021; Mitchell, 2011), and thereby fundamental to cellular homeostasis and bioenergetics. Consequently, membrane potential is used as an indicator for cell vitality (Lloyd and Hayes, 1995), which holds paramount importance across a wide range of biological and biomedical applications, encompassing the diagnosis of microbial infections, detection of proliferative cells, and examinations of synthetic-biology genetic circuits (Emerson et al., 2017; Hameed et al., 2018; Paternò et al., 2021; Riglar and Silver, 2018). Interestingly, recent studies also suggest that the resting plasma membrane potential may be related to cancer progression (Binggeli and Cameron, 1980; Yang and Brackenbury, 2013), indicating a fundamental link between the plasma membrane potential and cell vitality.

The plasma membrane potential can be influenced by an externally applied electric field (EFF), as described by the Schwan equation: Δ*ψ* = 1.5*rE* cos *θ*, where *r* is the radius of the cell, *E* is field strength and *θ* is the angle to the electric field (Marszalek et al., 1990). The change in membrane potential (Δ*ψ*) can then trigger the opening of voltage-gated ion channels (Rettinger et al., 2016). Since ion flux through channels follows the electrochemical gradient, the electrically induced membrane potential dynamics could be expected to be impacted by cell vitality (the capacity of cells to proliferate). In alignment with this idea, we previously showed proliferative-capacity-dependent dynamics in bacteria (Edwards et al., 2020; Stratford et al., 2019). In bacteria, the plasma membrane is a main site of ATP synthesis, which gives a possible explanation why bioenergetic status is linked to the electrically induced membrane potential dynamics. On the contrary, ATP synthesis does not happen at the plasma membrane in eukaryotes, instead it takes place in mitochondria. It is therefore unclear whether bioenergetic state of a cell alters the plasma membrane potential dynamics (PMD).Recently, a rise of fungal threats to human and ecosystem health and increased cases of antimicrobial resistance (AMR) has been observed (Fisher et al., 2012). The current protocol for anti-fungal susceptibility testing (AFST) is labour intensive and time-consuming, making the practical implementation difficult, posing a challenge for dealing with AMR. Electrically induced membrane dynamics of fungi cells could be a useful approach for antimicrobial susceptibility testing (AST) to evaluate novel and existing anti-fungal compounds and to optimise resistance surveillance. However, whether plasma-membrane potential response to electric stimuli may correspond to eukaryotic cell vitality is unknown.

In this study, we investigate the electrically induced membrane potential dynamics and its application for AST using the model organism *S. cerevisiae*. Our results suggest that the plasma membrane potential dynamics (PMD) shows hyperpolarisation in proliferative cells, but not in inhibited cells. Furthermore, mildly stressed cells showed an attenuated PMD response. Intriguingly, the dynamics correlate with the cellular growth rates, suggesting that electrically induced membrane potential dynamics can be used for bioelectrical AST (BeAST) to estimate the strength of growth inhibition.

## Results

Although the externally applied electric field (EEF) only directly interacts with the plasma membrane, the mitochondrial membrane plays a more central role in cell vitality. Therefore, we investigated the electrically induced dynamics of both plasma and mitochondrial membrane potential. To measure the plasma and mitochondrial membrane potential of *S. cerevisiae* cells, we used three fluorescent membrane-potential indicators: namely, bis-(1,3-dibutylbarbituric acid) trimethine oxonol (DiBAC4(3)), Tetramethylrhodamine methyl ester (TMRM) and Thioflavin T (ThT) (Figure 1A). DiBAC4(3) is an anionic dye that accumulates more in a depolarised plasma membrane; hence, its fluorescence intensity per cell increases as the plasma membrane potential becomes more positive, and decreases when it becomes more negative (Adams and Levin, 2012). TMRM is a cationic dye that is well established as a mitochondrial membrane-potential indicator that accumulates more in negatively polarised mitochondria (Scaduto and Grotyohann, 1999). ThT is a cationic benzothiazole dye that is most commonly used as a marker for amyloid fibrils but it is recently used also as a membrane-potential indicator in bacteria (Prindle et al., 2015; Sirec et al., 2019; Stratford et al., 2019), and also in mammalian cells (Skates et al., 2021).

**Figure 1.**
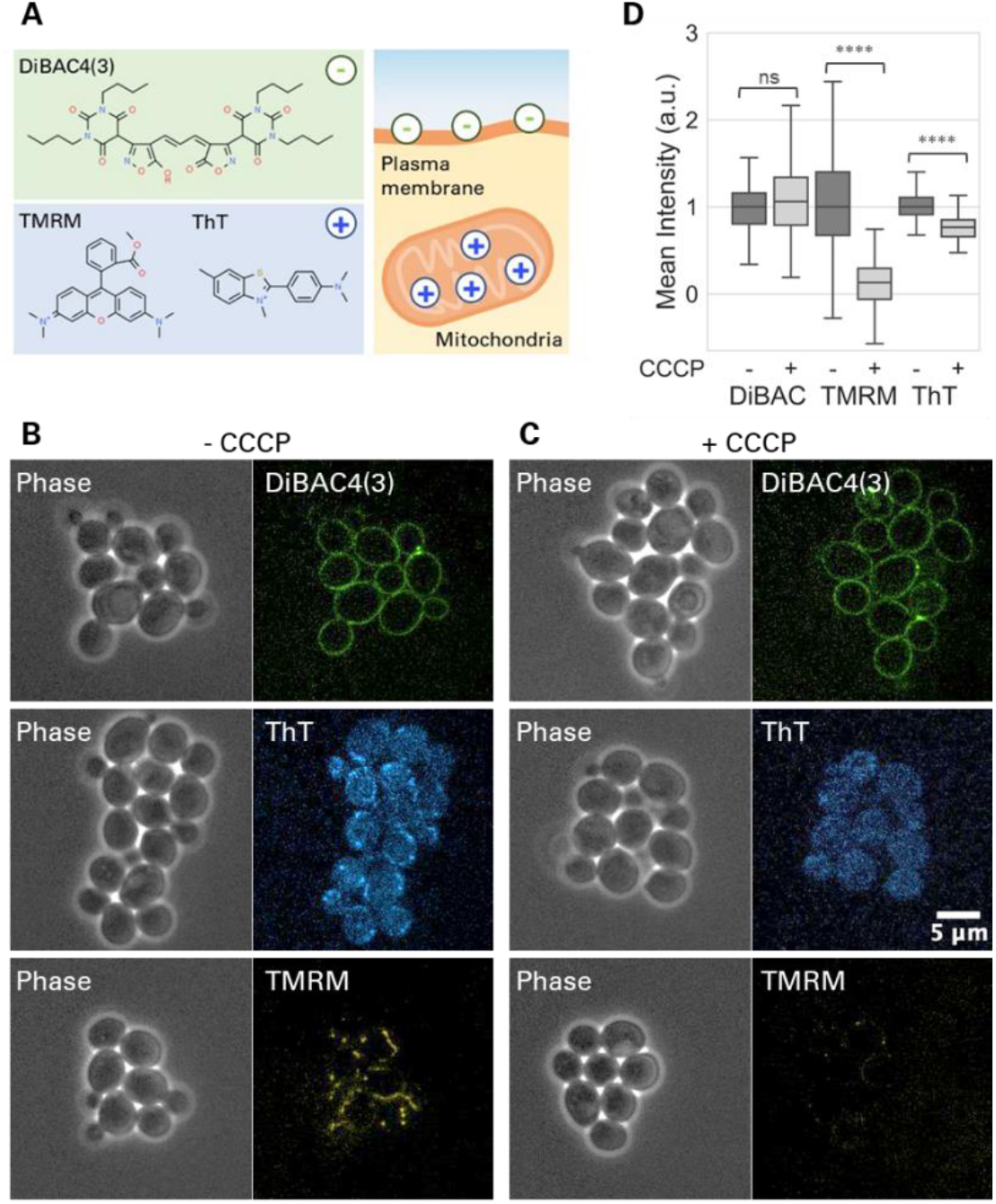
Plasma and mitochondrial membrane potential of *S. cerevisiae were monitored by fluorescece microscopy*. A) Illustrative diagram depicting the chemical structures of DiBAC4(3), TMRM and ThT. DiBAC4(3) is negatively charged and report on the plasma membrane potential. TMRM is positively charged and accumulate in the mitochondria. ThT is also positively charged and has been used to measure membrane potential. B, C) Microscopy images of *S. cerevisiae* cells shown in phase contrast (left) and fluorescence (right), B) without CCCP and C) with 10 μM CCCP. D) Box plot (median, 25^th^ percentile, 75^th^ percentile and range) showing the ratio change in the cellular fluorescence signals without and with CCCP treatment. Fluorescence intensities per cell were measured for individual cells. DiBAC4(3) (n>60 cells, p=0.34), TMRM (n>300 cells, p<0.01), and ThT (n> 300 cells, p<0.01) from 3 biological repeats were analysed.

To characterise the dyes in our experimental setup, we cultured *S. cerevisiae* cells on an agarose pad consisting of the Yeast Nitrogen Base (YNB) medium and supplemented with the membrane potential indicators. As expected, the fluorescence signals of DiBAC4(3) and TMRM are localised to the plasma membrane and in the mitochondria, respectively (Figure 1B). ThT showed mitochondrial accumulation similar to TMRM, although its signal appeared more diffused in the cytoplasm than TMRM (Figures 1B and S1). To test if this mitochondrial accumulation of ThT depends on the mitochondrial membrane potential, we treated cells with a mild level, at 10 μM, of a proton decoupler, Carbonyl Cyanide Chloro phenylhydrazone (CCCP) which depolarises the mitochondrial membrane but not the plasma membrane in eukaryotes (Skulachev, 1998). As expected, treatment with 10 μM CCCP resulted in the reduction of TMRM signal, but not DiBAC4(3) signal (Figures 1C, 1D). Note that this means that CCCP does not induce detectable changes in the plasma membrane potential. The fluorescence image of ThT showed a diffusive cytoplasmic pattern without a mitochondrial localisation signal (Figure 1C). This loss of mitochondrial localisation of the signal suggests that ThT accumulates in the mitochondria in a membrane-potential dependent manner. However, the diffusive signal of ThT was also visible. The quantification of the fluorescence signal confirmed that CCCP treatment only mildly reduced the ThT signal per cell (Figure 1D). These results suggest that ThT localises in mitochondria, but the mean ThT signal per cell is not highly sensitive to the mitochondrial membrane potential due to its diffusive cytoplasmic signal.

### The plasma membrane becomes more negative after electric shock

To examine the electrically induced plasma and mitochondrial membrane potential dynamics, YNB agarose pads carrying yeast cells were placed on a gold-coated electrode dish and used for stimulation experiment (Figure 2A). The dynamics of fluorescence signals before and after a 2.5-sec electric shock were monitored for 20 min by microscopy. The fold change in fluorescence signal over time (*I*(*t*)/*I*_0_) was used to quantify the response dynamics.

**Figure 2.**
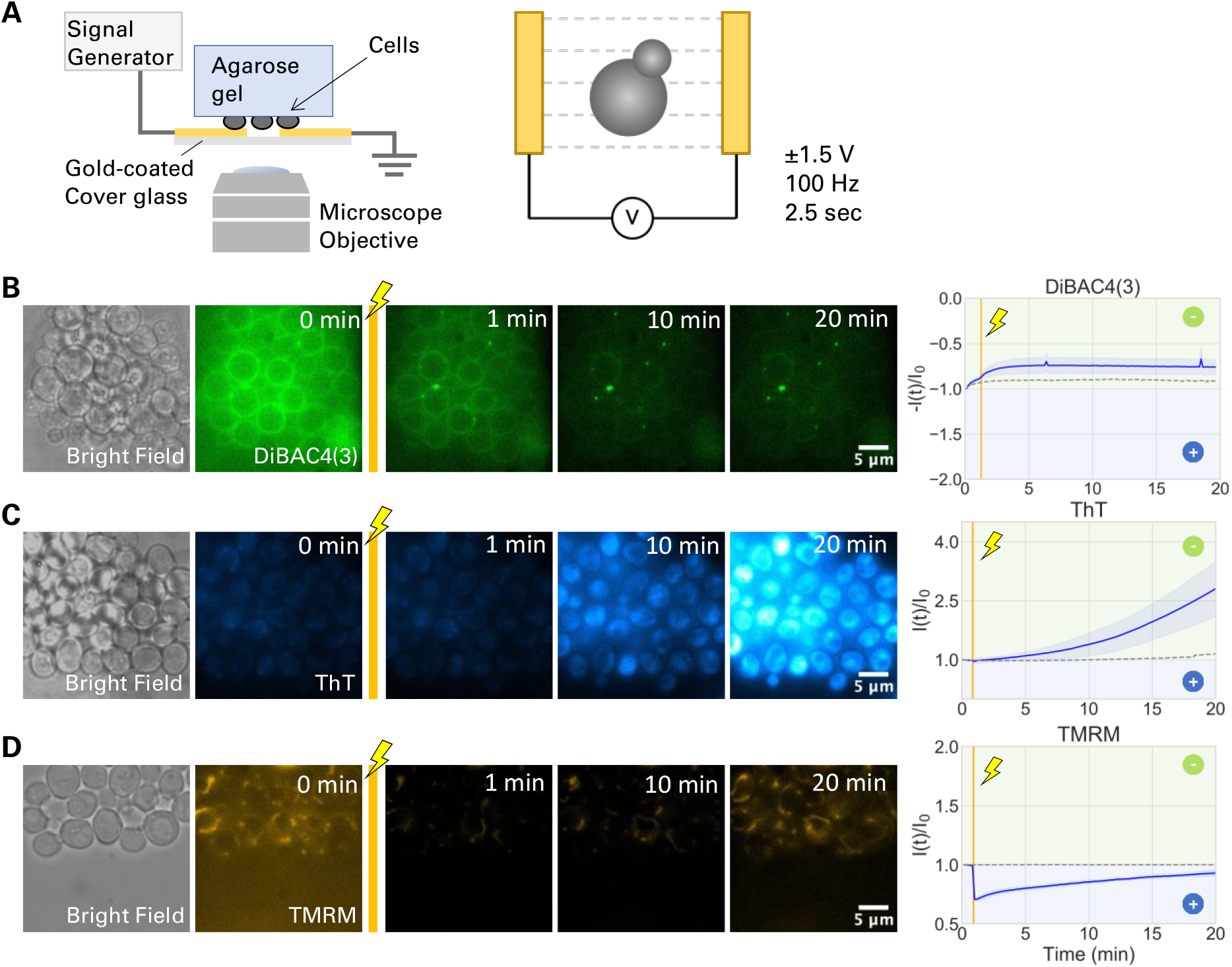
Electric stimulation induces plasma membrane hyperpolarisation. A) Schematic illustration of the experimental setup. Yeast cells on an agarose pad are placed facing down onto gold electrodes on a coverglass-bottom dish, connected to a signal generator and mounted to a inverted fluorescence microscope. A ±1.5 V, 100Hz stimulus is applied for 2.5 seconds. B-D) Proliferative cells exhibit a hyperpolarisation in response to EEF. B-D) Bright field images and film-strips of fluorescence microscopy images before and after electrical stimulation. Graphs showing the fold change in fluorescence signal over time (blue solid line). Shaded blue shows standard deviation. Dashed lines are the condition without electrical stimulation. The color in the background in the graphs show the direction of change in membrane potential: limegreen shows membrane potential going further negative, blue shows it going more positive. Gold vertical line indicates 2.5-sec electrical stimulation. B) DiBAC4(3) signal shows a quick and stable hyperpolarisation, depicted by decrease in fluorescence intensity. C) ThT signals show a gradual hyperpolarisation, with 3 fold increase in fluorescence intensity. D) TMRM signals shows an instant depolarisation in response to EEF, followed by a gradual return to the initial resting membrane potential.

Following the 2.5-sec stimulation, DiBAC4(3) signal decreased by ∼25% over the period of 20 min, suggesting hyperpolarisation of the plasma membrane (Figure 2B and Movie 1). The hyperpolarisation following electric stimulation was also observed with ThT, which showed ∼3x rise in the signal (Figure 2C and Movie 2). The ThT signals after electric stimulation were more diffusive in the cytoplasm, which implied less mitochondrial accumulation due to mitochondrial membrane depolarisation. TMRM signal showed a rapid decline after an electric stimulation, suggesting mitochondrial depolarisation (Figure 2D and Movie 3). In the absence of electric stimuli, no significant change in the fluorescence signal was observed over the course of the 20-min time-lapse experiments in absence of electrical stimulation (Figure S2). The dynamics of individual cells for all three dyes are presented in Figure S3. These results suggest that the electric stimulation hyperpolarises the plasma membrane while it depolarises the mitochondrial membrane.

### Cell vitality is required for electrically induced hyperpolarisation

To examine if the electrically induced membrane-potential dynamics relates to cell vitality, we used UV-Violet light to inhibit cells. Specifically, the cells within the centre region of the field of view were exposed to UV light for 1 min whereas cells in other locations were not (Figure 3A and B). This experimental design enabled us to directly compare UV-exposed and non-exposed cells within the same field of view while ensuring that they received identical electric shock.

**Figure 3.**
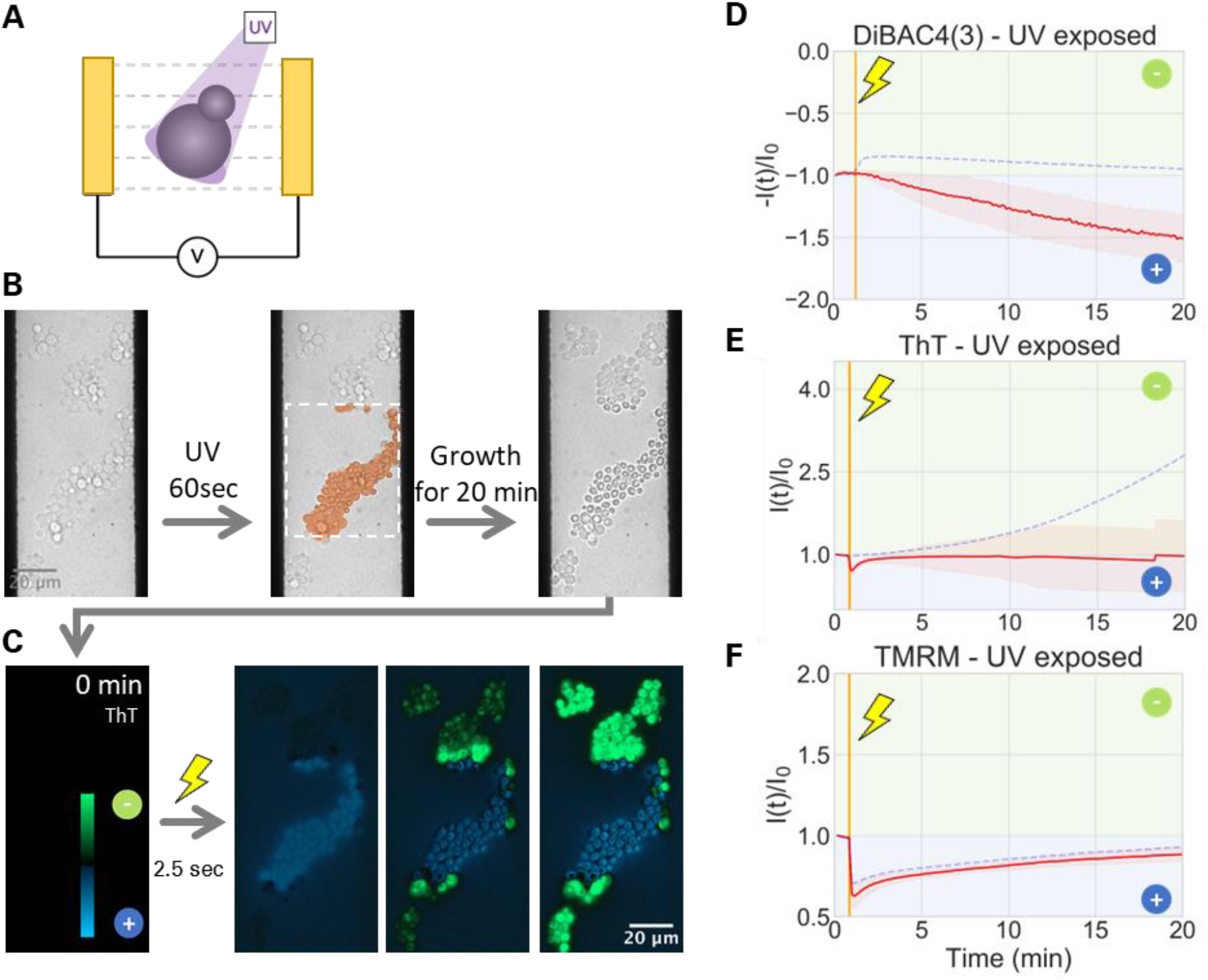
UV-irradiated cells do not produce a hyperpolarisation response to EEF. A) Schematic illustration of experimental design. B) Flowchart diagram showing the experimental design. Bright field images show *S. cerevisiae* cells within the electrode gap. UV-Violet irradiation for 60-sec is applied only to the centre of the field of view (highlighted by the dashed rectangle). The growth inhibition by the irradiation is verified by further incubation for 20 min and brighlight field time-lapse microscopy (see also Figures S3 and S4). Cells are then exposed to EEF for 2.5 sec. C) The ratio change of ThT signal in log scale is calculated from microscopy (log(*I*(*t*)/*I*_0_)) and shown in a green-black-blue pallete . Green shows hyperpolarisation and blue shows depolarisation. The images correspond to panel B. D-F) Graphs showing the fold change in fluorescence intensities of UV-V irradiated cells (solid red line) and unirradiated cells (dashed blue). Shaded red shows standard deviation. For the negative control without electrical stimulation, see Fig. S2. The color in the background in the graphs show the direction of change in membrane potential: limegreen shows membrane potential going further negative, blue shows it going more positive. Gold vertical line indicates 2.5-sec electrical stimulation. D, E) DiBAC4(3) and ThT show a lack of hyperpolarisation response for UV-V irradiate cells (solid line), but not for non-irradiate cells (dashed line). F) TMRM shows an instant depolarisation followed by a gradual recovery, regardless of UV-V irradiation.

We first confirmed by bright-field time-lapse microscopy that UV-exposed cells stopped growing (Figures 3B, S4 and S5). The exposed cells also appeared severely damaged in brightfield images and fluorescence images (Figures S5 and S6). With all three fluorescence dyes that we tested, fluorescence signals were brighter and diffusive in the cytoplasm (Figure S6). It must be noted that both anionic and cationic dyes increased their signals diffusively in the cytoplasm. This suggests that this rise in intensity is not due to membrane potential changes, and their application for indicating membrane potential is limited with UV-stressed cells. With this potential limitation in mind, UV-exposed cells were stimulated by an electric field and the fluorescence signals were monitored for another 20 min (Figure 3). We analysed the fold change in fluorescence intensities over time (*I*(*t*)/*I*_0_). The response dynamics of TMRM signal was indifferent between UV-exposed and non-exposed cells and showed a transient drop in fluorescence signal (Figures 3F and S7). This could be due to the fact that the mitochondrial membrane is insulated by the plasma membrane and would not be directly exposed to EEF.

The anionic dye DiBAC4(3) showed a ∼50% rise in exposed cells (Figures 3D and S7), which is in a clear contrast to non-exposed cells (Figure 3D, dashed line). Intriguingly, electrically induced dynamics of ThT signal was also different between non-exposed and exposed cells. In exposed cells, ThT signal showed a slight transient decrease followed by a largely flat signal (Figures 3E and S7), while non-exposed cells showed a gradual rise of ThT signal (Figure 3E, dashed line). The ratio changes in ThT fluorescence are visualised in Figure 3C. In these images, a small number of cells within the irradiated region showed ThT signal increase near an electrode (Figure 3C and Movie 4). However, closely examining the pre-shock bright-field images, we found that these cells entered the region during the incubation period –hence not exposed to UV light. Putting together the dynamics of an anionic dye, DiBAC4(3), and a cationic dye, ThT, our results strongly suggest that the UV-inhibited yeast cells are unable to produce a hyperpolarisation response to electric stimulation. In other words, cells need to be vital to produce an electrically induced hyperpolarisation response.

To further examine if stressed cells show altered response dynamics of membrane potential, we performed the assay with the cells treated with ethanol. Ethanol-treated cells showed plasma membrane depolarisation upon electric stimulation (Figure S8). These results suggest that membrane potential response to electric stimulation can be used for rapid antimicrobial susceptibility testing (AST), which we name as bioelectrical AST (BeAST).

### The FHN mathematical model recapitulates vitality-dependent hyperpolarisation

The cell-vitality-dependent hyperpolarisation prompted a question about how mildly stressed, but not fully inhibited, cells react to an electric shock. In particular, we wonder if the response dynamics is gradual or binary. To explore this question, we employed the FitzHugh-Nagumo (FHN) model, a phenomenological mathematical model for neural activity (FitzHugh, 1961). The FHN model is an ODE system with a cubic nonlinear function for membrane potential (V) and a linear function for a recovery variable (W) which represents slow negative feedback. In this model, similar to our previous study on bacteria electrophysiology (Stratford et al., 2019), vital cells accumulate high level of potassium inside the cell, which can be released upon electric shock. On the other hand, because maintaining the membrane potential is a major energetic burden (Milo and Phillips, 2015), non-vital cells are unable to accumulate high intracellular potassium, leading to the absence of hyperpolarisation.

We numerically simulated the dynamics of the plasma membrane potential after an electric stimulation with varying levels of cell vitality (*k*). This parameter *k* has two effects on the model behaviour. First, it shifts the equilibrium point, thus the resting membrane potential levels. Second, it determines the time scales of membrane potential dynamics, relative to the time scale of recovery variable (W). More specifically, with smaller *k* (i.e. less cell vitality), membrane potential dynamics becomes more gradual. As a result, smaller *k* gives rise in a weaker hyperpolarisation response. This dependency on *k* illustrated by simulation results indicates a smaller hyperpolarisation response with lower cell vitality (Figure 4A).

**Figure 4.**
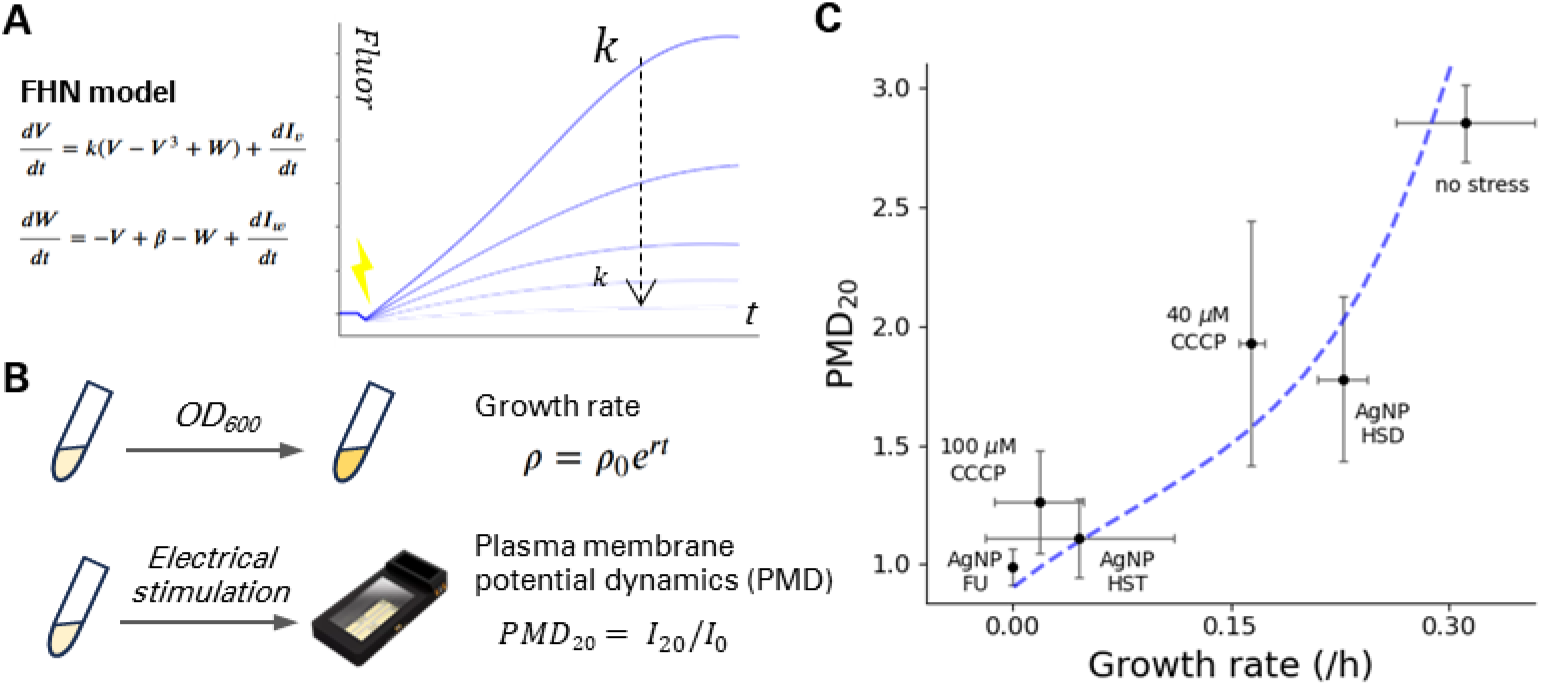
Electrically induced membrane potentail change correlates with growth rates. A) The FHN model is used to simualte the electrically induced membrane potentia dynamics. Simulations are performed with various values for *k* (0.1, 0.3, 0.5, 0.7, 0.9) and the changes in membrane potential (Δ*ψ*) over time are plotted. B) Cells in various conditions are cultured to estimate the growth rate (*r*) and the plasma membrane-potential dynamics (PMD) of electrically induced membrane potential change at 20 min (PMD_20_). C) The relationship between growth rate and electrically induced membrane potential dynamics. To calcualte the growth rates, OD_600_ was measured in liquid culture. PMD_20_ was calculated from fluorescece time-lapse microscopy images. Text in the graph shows stressors. Spearman r value = 0.89, p value = 0.019. The light blue dashed line shows the FHN simulation with various k values.

### Milder stress reduces hyperpolarisation level

To test this conjecture from the model, we performed plate-reader assays to measure OD_600_ for 22 hours and calculated the growth rate (r) for the cells treated with and without stressors (Figure 4B). We also performed BeAST with the cells treated with stress agents (Figure 4B). ThT was used for this experiment because it showed a greater fold-change difference between UV-damaged and untreated cells than DiBAC4(3) (Figure 3E). The plasma membrane potential dynamics (PMD) before and after electric shock was calculated to quantify the magnitude of hyperpolarisation.

We used the proton decoupler CCCP as a known chemical stressor. Note that CCCP does not cause a detectable change in the plasma membrane potential (Figure 1). Culturing cells with varying concentrations of CCCP between 0 and 100 μM, we observed a dosedependent growth inhibition of CCCP (Figure S9). The growth rates were reduced by ∼50% and ∼90% with the addition of 40 and 100 μM CCCP, respectively.

For the electrical stimulation experiment, cells were exposed to 40 and 100 μM CCCP for 1 hour and used for time-lapse fluorescence microscopy. The membrane potential dynamics before and after electrical stimulation was monitored using ThT (Figure S9) and PMD_20_ was calculated. Intriguingly, PMD_20_ was lowered by CCCP in a dose-dependent manner. Plotting PMD_20_ against the growth rate with and without CCCP, we find a correlation between growth rate and PMD_20_ (Figure 4C). This result suggests that membrane potential dynamics can be used for assessing antifungal activities.

### BeAST can be used for examining antifungal activities of silver nanoparticles

The above findings suggest that BeAST can accelerate the assessments of novel antifungals as it can be completed in a few hours while a growth assay takes overnight. We explored this possibility using biogenic silver nanoparticles (bioAgNPs), promising alternative to antibiotics. We used the methods that can reproducibly produce stable AgNPs (Ballottin et al., 2017; de Barros et al., 2018; Stanisic et al., 2022). The methods are also economically and ecologically beneficial as they can utilise orange wastes and do not produce toxic waste in the process. Following a few non-time-consuming steps, we obtained AgNP-HST and AgNP-HSD using orange fruit flavonoids. We also produced bioAgNP using the filamentous fungus *Fusarium oxysporum* (Stanisic et al., 2022). The basic properties of bioAgNPs were characterised by electron microscopy, spectroscopy and dynamics light scattering (Figures S13-S15, Table S1). *S. cerevisiae* cells were incubated with AgNPs for 1 hour in liquid culture and further incubated on pads for <3 hours before stimulated by an electric shock. The PMD_20_ was then measured using fluorescence time-lapse microscopy images. AgNP-FU-treated cells were reminiscent of the UV-exposed cells and showed a reduced PMD_20_ (Figure S10). Amongst two orange NPs, HST showed a greater reduction in PMD_20_ than HSD (Figure S11, S12). We also performed overnight plate-reader assay to determine the AgNP’s effects on the growth rates. Intriguingly, the plate-reader assay showed that HST and FU nanoparticles are more effective in inhibiting growth than HSD, which was consistent with PMD_20_ (Figure 4C). Therefore, this result suggests that BeAST can be used for examining antifungal activities based on membrane potential response dynamics to electric stimulation.

## Discussion

This study presents bioelectrical AST (BeAST) based on the finding the electrically induced dynamics of the plasma membrane potential correlates with the inhibition of growth rates. We observed a correlation between growth rates and hyperpolarisation, suggesting that BeAST can estimate the magnitude of stress. Strikingly, this relationship appears to be independent from molecular modes of action of antimicrobials.

We show that an electric stimulation hyperpolarises the plasma membrane. This conclusion is supported by the measurements with two membrane-potential indicators: anionic DiBAC4(3) and cationic ThT. These results indicate that electrophysiological dynamics in response to EEF does not depend on a specific dye. Furthermore, it suggests that this electrophysiological response dynamics may be an evolutionary conserved feature, since the prokaryotic organisms *Bacillus subtilis* and *Escherichia coli*, and the eukaryotic model *Saccharomyces cerevisiae*, show similar membrane potential dynamics in response to an electrical shock (Stratford et al., 2019). The exact mechanism by which an electric stimulation triggers such an event requires further investigation. For instance, it is not clear whether EEF directly or indirectly impacts cellular membrane and membrane proteins. An important next step for microbial electrophysiology is to characterise the mechanism and explore possible explanations for why such a feature may be conserved.

We believe that BeAST has many potential applications in both industrial and laboratory settings. When applied to cellular populations, such an assay can be used to determine viability, defined as the number of vital cells. *S. cerevisiae* is an organism with significant applicability in biotechnological processes, which range from brewing and baking to the production of pharmaceutical molecules and expression of heterologous proteins (Buckholz and Gleeson, 1991; Kingsman et al., 1985). However, yeast can also be a concerning contaminant in industrial settings, with 9% of non-food product recalls being caused by yeast and mould (Cunningham-Oakes et al., 2019). Thus, rapid assessment of vitality and viability to assure either efficient bioprocesses in food and biotechnology industries, or efficient elimination of contaminants, is a pressing need. The traditional methods for monitoring yeast viability rely on growth analysis methods including colony forming units (CFU) and spotting tests, or on live/dead cell staining methods including methylene blue and phloxine B (Kwolek-Mirek and Zadrag-Tecza, 2014). These methods, however, are not suited for measuring vitality. Vitality measurements generally rely on monitoring cellular growth or metabolic activities. While these methods are widely used and well established, conducting and analysing them take several days or require complex calibrations. Flow cytometry is a sophisticated technology that can provide rapid results of live cell detection; however, it requires technical expertise and its ability to estimate the growth rate may be limited. Our work suggests that electrical stimulation can shorten the time it takes to do such an assay. Additionally, this approach could accelerate the screening of anti-fungal compounds and potential therapeutic agents for fungal infections. It could also be useful for basic research using *S. cerevisiae* where measuring cell growth rates is important. For example, the growth rate and electrophysiology properties of *S. cerevisiae* have been shown to correlate to ageing (De Souza-Guerreiro and Asally, 2021; Haupt et al., 2014). A future study shall examine the relationships between PMD and CCCP concentrations more granularly and explore different types of stressors to quantitatively characterise the relationship between stress and plasma membrane potential dynamics. It is yet to be seen if BeAST can be applied with clinically relevant fungal species and strains. If they do, estimating the growth inhibition levels by BeAST may shorten the time it takes for assays and could uncover the roles that bioelectricity plays in a plethora of cellular processes. Thus, BeAST has a potential to aid the diagnosis of fungal infections and development of antifungals.

## Materials and Methods

### Strain and growth condition

*Saccharomyces cerevisiae* (haploid, W303 background) was cultivated in yeast nitrogen base (YNB) (Foremedium) with Complete Supplement Mix (CSM) (Foremedium) liquid media supplemented with 2% glucose (wt/vol) and on yeast extract peptone dextrose (YEPD) agar (1.5% [wt/vol]). For time-lapse and electrical stimulation experiments, a single colony of *S. cerevisiae* was inoculated into YNB + 2% glucose media and cultured at 30°C in a shaking incubator overnight. In the following morning, cells were diluted into YNB + 2% glucose to OD_600_= 0.15 - 0.2 and allowed it to grow at 30°C, shaking until reaching OD_600_ = 0.6. When specified so in figure legend, Carbonyl Cyanide Chlorophenylhydrazone (CCCP) was added to the liquid culture to the final concentration of either 10, 40 or 100 μM in the last 1 hour of incubation. Cells were then inoculated onto YNB 2% glucose supplemented with low-melting-point agarose (1.5% [wt/vol]) pads with membrane potential dye. Pads were prepared as described previously (Young et al., 2011) by melting agarose (Fisher bioreagents – Low melting point agarose) into YNB 2% glucose liquid media, then pouring 1 mL onto 22 mm x 22 mm glass slip and covering with another glass slip to it to solidify. Agarose was cut into 5 mm x 5 mm pads. 3∼5 μL of culture was inoculated on individual pads that were then placed onto a willco dish for general time-lapse experiments or placed onto a gold-coated glass-bottom dish for electrical stimulation experiments (Edwards et al., 2020; Stratford et al., 2019).

### General time-lapse experiments

Yeast membrane potential dynamics was observed using a Nikon TI-E Eclipse motorized wide-field epifluorescence microscope equipped with an EM-CCD camera (Andor DU-888). Operation of the microscope was performed using NIS Elements (Nikon) and temperature control was obtained by an incubation chamber (Okolab Bold Line Cage Incubator). Before time-lapse experiments, chamber temperature was set at 30°C for at least an hour; then samples were placed into the chamber a further hour before starting the experiment. Time-lapse experiments were performed using 100x objective lens (N.A. 1.45, Nikon). Cell growth was recorded using phase contrast. Membrane potential dynamics was monitored by the combination of the excitation laser and emission filter adequate for the dye used in that specific experiment. ThT fluorescence was detected using a 405 nm excitation laser and a QUAD emission filter. TMRM fluorescence was detected using a 561 nm excitation laser and a QUAD emission filter. DiBAC fluorescence was detected using a 488 nm excitation laser and a GFP-FITC (525/50) emission filter.

### Electrical stimulation experiments

Electrical stimulation experiments were performed as described previously (Edwards et al., 2020; Stratford et al., 2019). Briefly, stimulation was performed by applying an alternating current (AC) signal [0.1 kHz; 3 V peak-to-peak (−1.5 ∼ +1.5 V)] to individual electrodes of the gold-coated glass-bottom dish using CytePulse (Cytecom Ltd). Cell growth was monitored prior to electrical stimulation using an inverted epifluorescence microscope, DMi8 (Leica Microsystems), operated by MetaMorph (Molecular Devices) and equipped with a scientific CMOS camera ORCA-Flash 4.0 v2 (Hamamatsu Photonics). All experiments were conducted at 30°C, maintained by an i8 incubation chamber (Pecon). Samples were placed into the chamber for 1-2 hour for thermal equilibration. Observations were performed using a 100× objective lens (N.A. 1.3, HCX PL FLUOTAR; Leica). Growth was observed with bright field and membrane potential dynamics was monitored using ThT, TMRM or DiBAC4(3). ThT fluorescence was detected using a 438/24 nm excitation filter, a 483/32 emission filter and a 458 dichroic mirror (Semrock). TMRM was detected using a 554/23 excitation filter, a 609/54 emission filter and a 573 dichroic mirror (Semrock). DiBAC4(3) was detected using a 500/20 excitation filter and a 578/105 emission filter. Prior to electrical stimulation, cell growth was recorded for 1 hour. For UV light exposure experiments, cells were exposed to UV-Violet (UV-V) light for 1 minute, using the DMi8 inverted microscope (Leica Microsystems) and the LED light source, SOLA SM II Light Engine (Lumencor) equipped with an excitation filter of 400/16nm. The area exposed to UV light was controlled by the field diaphragm of the microscope.

### Biogenic Silver Nanoparticles

Biogenic silver nanoparticles were prepared using the protocol described by Stanisic et *al*. (Stanisic et al., 2022), using flavonoids from oranges - hesperetin (HST) and hesperedin (HSD) - as reducing and stabilising agents. Hesperidin (HSD) and its aglycone, hesperetin (HST), are two flavonoids from citrus species that have various biological properties and take part in a biogenic silver nanoparticle (AgNP) synthesis in the same way, using flavanone moiety in an oxidation/reduction reaction. Along the synthesis, HSD and HST stabilize the AgNPs by interaction with the AgNP surfaces. Therefore, it was expected that AgNP-HSD and AgNP-HST would show very similar properties and activities. The only structural difference is the disaccharide rutinose moiety in HSD as illustrated below. The synthesised AgNP-HSD and AgNP-HST were spherical in shape (de Barros et al., 2018), exhibited low zeta potentials (around -35 meV), polydispersity around 0.4, and 25-30 nm in diameters. The unique difference between them was the sugar moiety (O-rutinoside) on HSD. We have fully characterized AgNPs using dynamic light scattering, surface charge measurements, and transmission or cryogenic electron microscopy. See also supplementary information. Fungal biogenic silver nanoparticles (AgNP-FU) were prepared according to the protocol described by Ballotin et al. (2017) (33) using the fungal filtrate prepared from the Fusarium oxysporum biomass. The synthesised AgNP-FU were spherical, exhibited low zeta potential (-34.5 ± 5.4 mV), acceptable polydispersity, and 25 nm in diameter. Using dynamic light scattering, surface charge measurements, and cryogenic transmission electronic microscopy, AgNP-FU particles were fully characterised as shown in the Supplementary Information.

### FttzHugh-Nagumo model

The electrically induced membrane potential dynamics was simulated using the FitzHugh-Nagumo (FHN) neuron model, similarly to a previous work (Stratford et al., 2019). The membrane potential *Vm* and the recovery variable *W* were considered by the following equations:

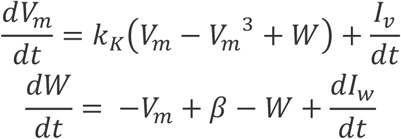

where *I*_*v,w*_ is the externally applied electrical field (EEF) and *k*_*K*_ represents cell vitality. *B* was defined as:

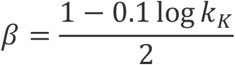

The equations were solved in Python using a scipy ode solver (scipy.integrate.odeint). EEF was applied to the equilibrium state. To analyse the impact of cellular states, the simulations were conducted for *k*_*K*_ between 0.1 and 1 and the ratio changes in V were recorded. The fluorescence intensity (I) was then calculated as *I* = *e*^*αV*^. The ratio change in I was plotted. The growth rate was assumed to be linearly proportional to *k*_*K*_.

## Acknowledgements

We thank Drs Jintao Liu, Orkun Soyer, Fabio Dos Santos, Magdalena Karlikowska for the discussion and the comments to the manuscript. We thank Conor Edwards for his help with experiments and discussions, and the members of Asally lab and the Warwick Bio-Electrical Engineering Innovation Hub for discussions. We acknowledge the funding by BBSRC Impact Acceleration Award (BB/S506783/1), Flexible Talent Mobility Account (BB/S507982/1), the University of Warwick Research Development Fund (RDF) Strategic Award, the Royal Society International Exchange grant (IES\R3\223259) and EPSRC/BBSRC synthetic biology centra grant (BB/M017982/1). We also acknowledge the funding by São Paulo Research Foundation (FAPESP, Grant #2014/50867-3, #2023/02338). This research used facilities of the Brazilian Centre for Research in Energy and Materials (CNPEM), a private non-profit organisation under the supervision of the Brazilian Ministry for Science, Technology, and Innovations (MCTI). The TEM, LNNano staff is acknowledged for their assistance during the experiments (TEM-C1-20210476 and TEM-27348 - CryoEM).

