## Supplementary material for "Membrane potential dynamics unveil the promise of Bioelectrical Antimicrobial Susceptibility Testing (BeAST) for anti-fungal screening": SI

**This PDF file includes:**

Supplementary Figures S1 to S15
Supplementary Table S1
Supplementary Movies S1 to S4
Supplementary Materials and Methods

**Other Supplementary Materials for this manuscript include the following:**

Movies S1 to S4

### Supplementary Figures

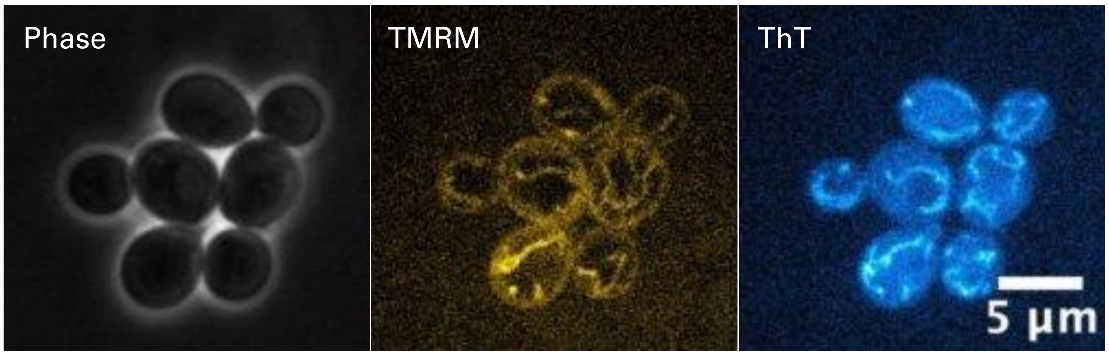


**Fig. S1**. Co-staining of Saccharomyces cerevisiae cells with TMRM and ThT. Images show co-localization of both dyes in the mitochondria, suggesting ThT accumulates in this organelle, reporting on its membrane potential.

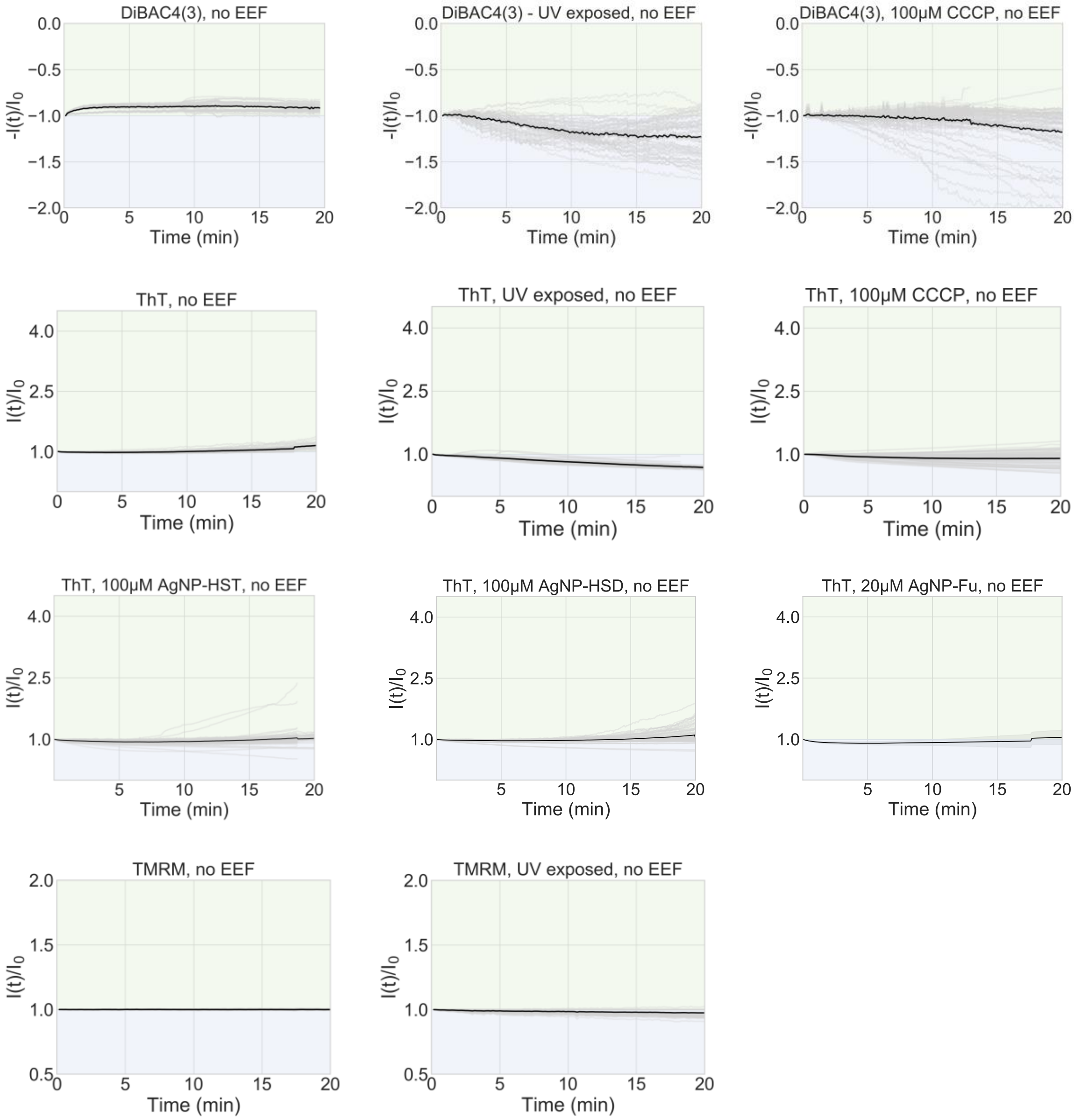


**Fig. S2**. Cell response without externally applied electrical field. Light grey lines are single-cell time traces. Black line is the mean of single-cell time traces.

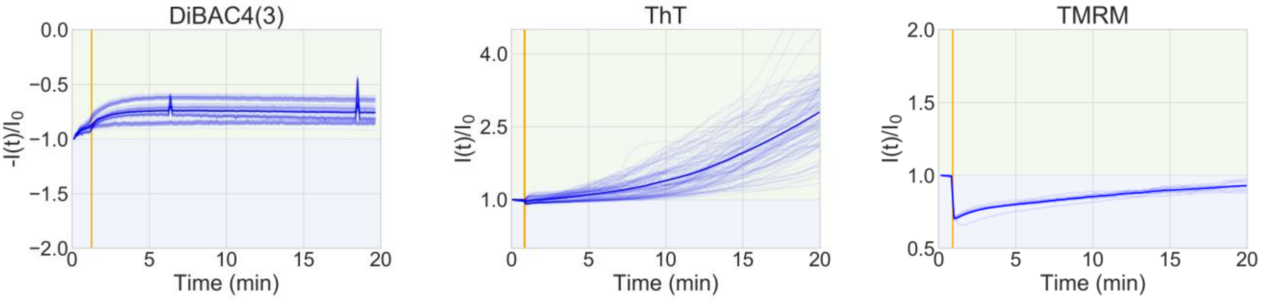


**Fig. S3.** Single-cell time traces of proliferative yeast cells exposed to EEF, represented by thinner blue lines. Thicker lines represent the mean of single-cell traces

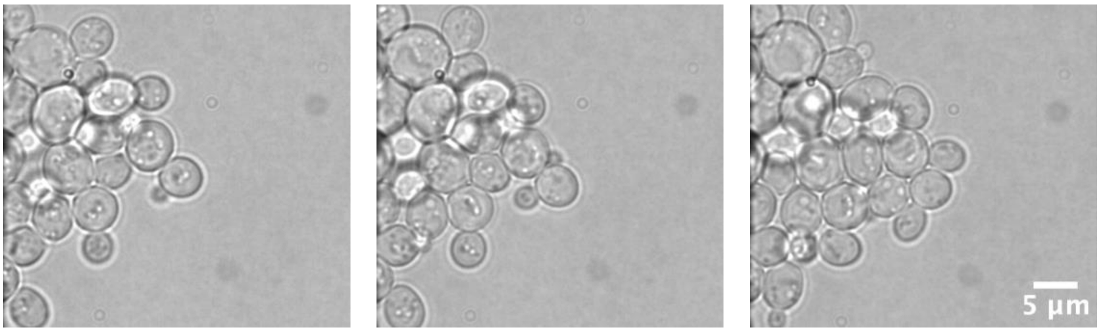


**Fig. S4**. Healthy cells were observed for one hour to assess their proliferative capacity. Bright field images depicting cellular growth on the electrode dish.

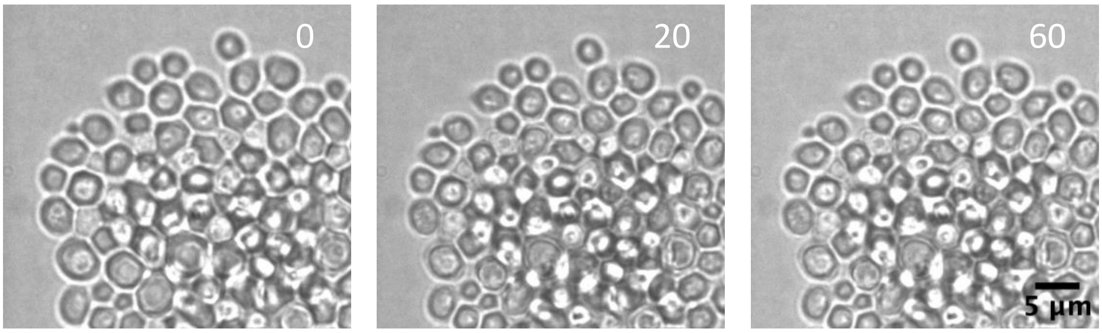


**Fig. S5**. UV-V irradiated cells were observed for one hour to assess growth inhibition. Bright field images showing successful inhibition of cell division.

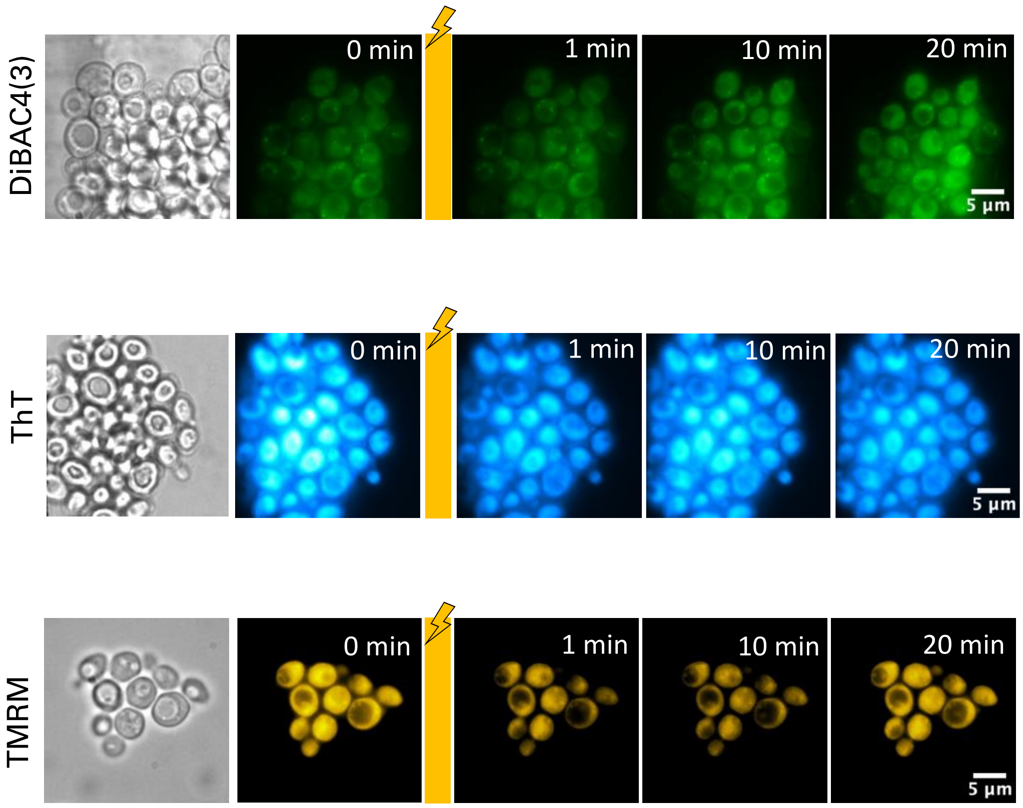


**Fig. S6.** UV-V irradiated cells were observed for one hour to assess growth inhibition. Bright field images showing successful inhibition of cell division.

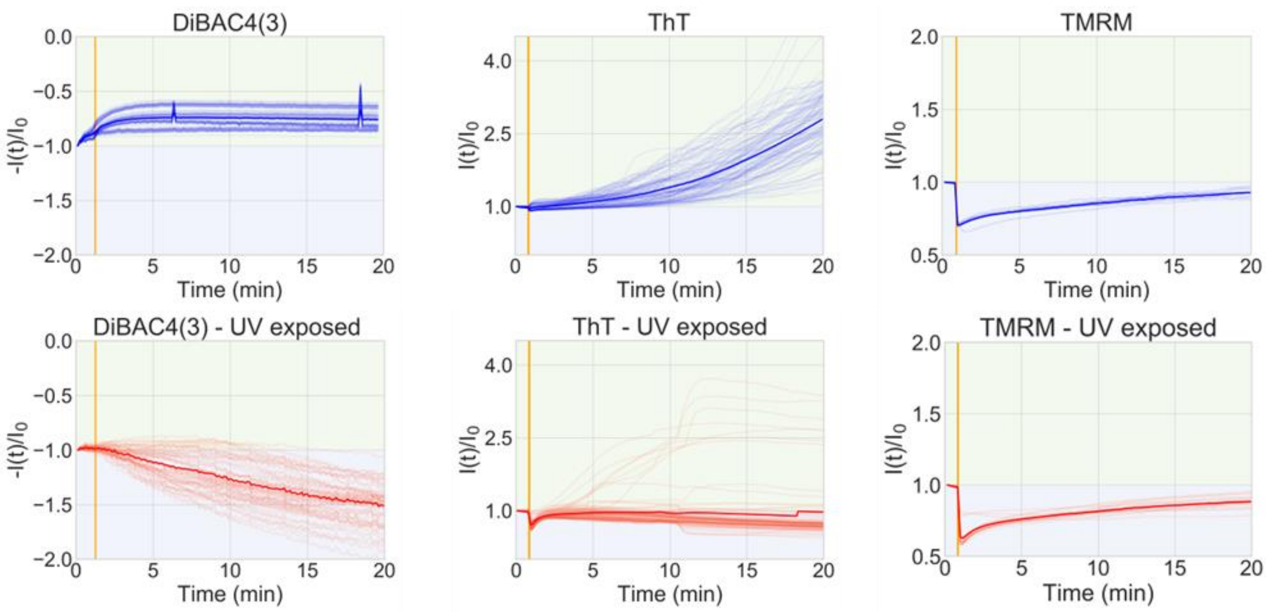


**Fig. S7.** Single-cell time traces of proliferative or UV-inhibited cells exposed to EEF, represented by thinner lines. Blue represents proliferative cells and red represents inhibited cells. Thicker lines represent the mean of single-cell traces.

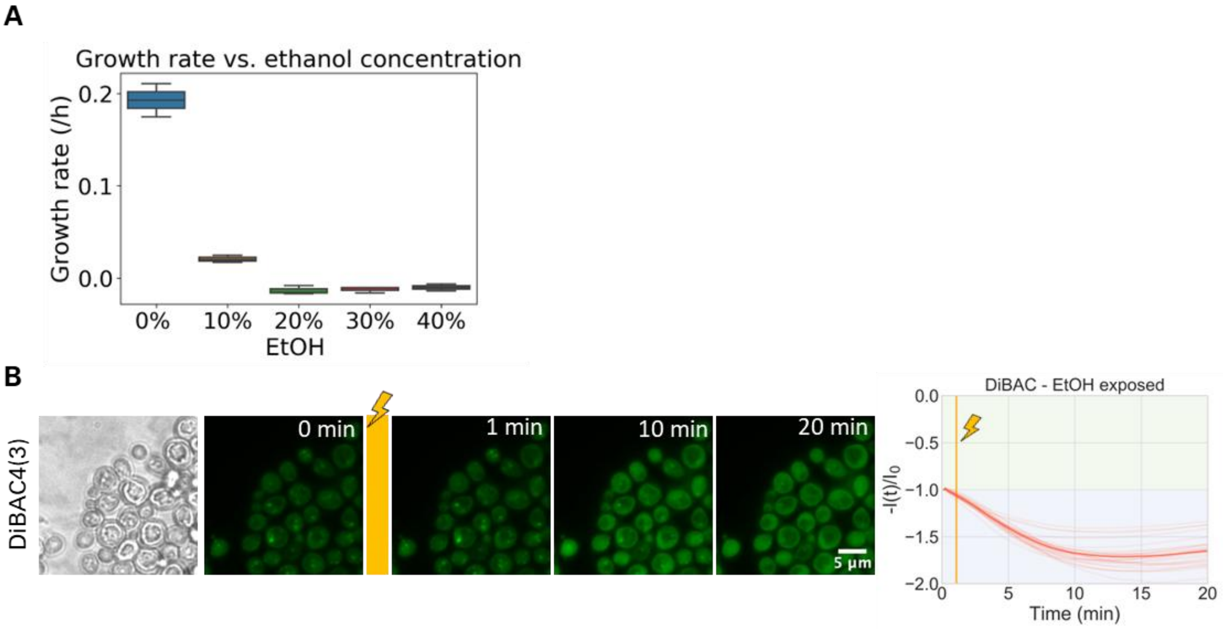


**Fig. S8.** Ethanol 30% v/v inhibits growth and induces depolarization upon EEF. (A) Increasing concentrations of ethanol inhibits growth, with 30 and 40% v/v completely impairing proliferation. (B) Cells stained with DiBAC4(3) show a gradual depolarization, depicted by a 1.5 fold increase in fluorescence intensity.

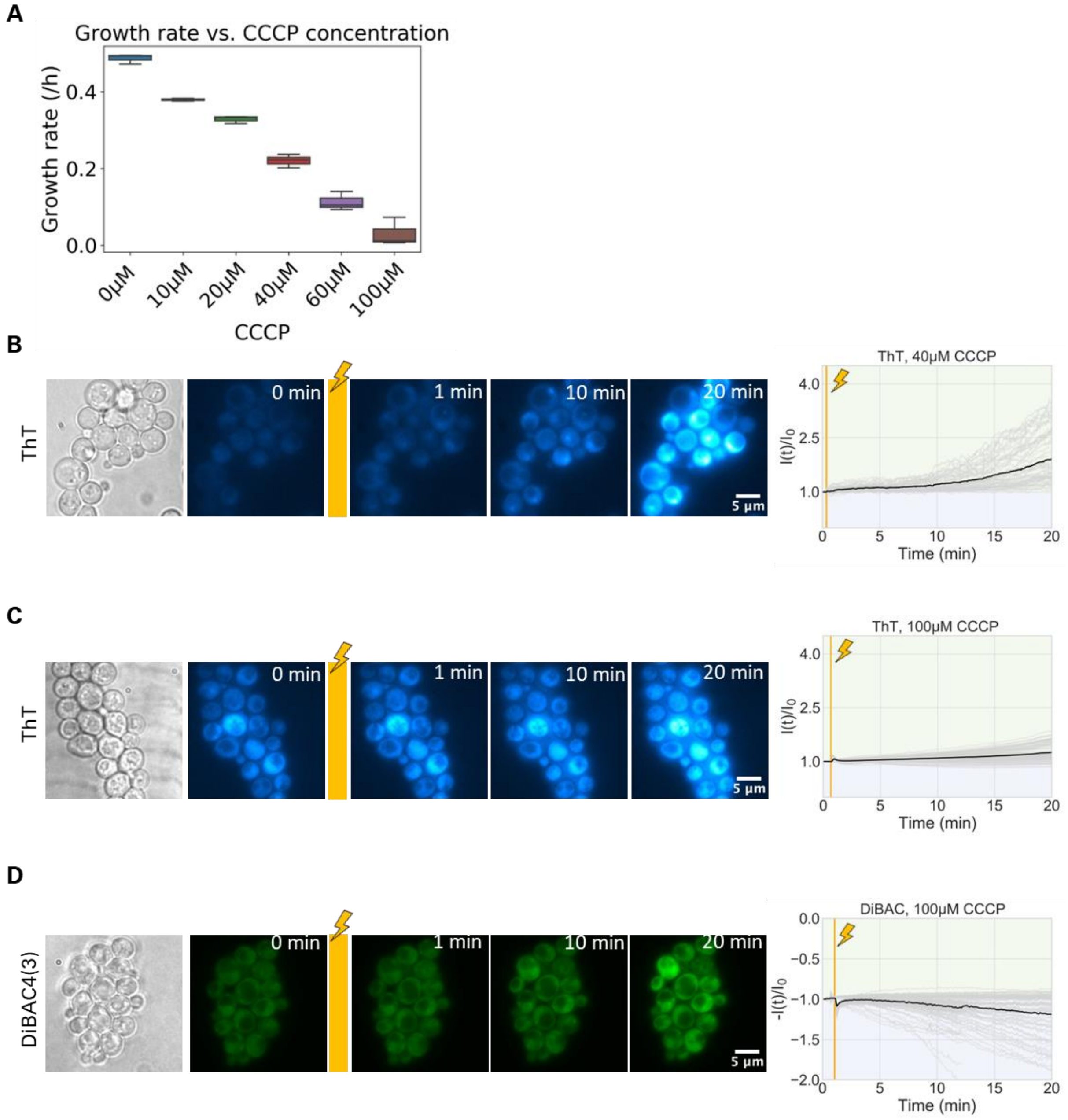


**Fig. S9.** The protonophore CCCP delays growth in a concentration-dependent manner. (B) Cells stained with DiBAC4(3) exposed to 100µM CCCP show a gradual depolarization, less intense than cells irradiated with UV-V or exposed to EtOH 30%v/v. The same trend is observed with ThT: (C) Cells exposed to 40µM CCCP exhibited delayed growth, with response to EEF being a intermediate, 2 fold hyperpolarization. (D) Cells exposed to 100µM CCCP show a discrete hyperpolarization, demonstrating that the membrane potential dynamics in response to an EEF repots not only cell viability, but also on cell vitality.

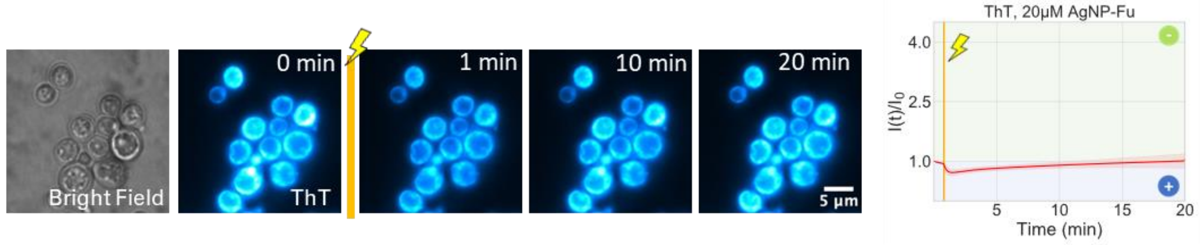


**Fig. S10.** AgNP-Fu exerts a strong inhibitory affect in fungal cells, which in turn depolarize immediately after the shock and maintain a relatively stable dynamics after EEF stimulation. Shaded areas represent the standard deviation, while the lines represent the mean.

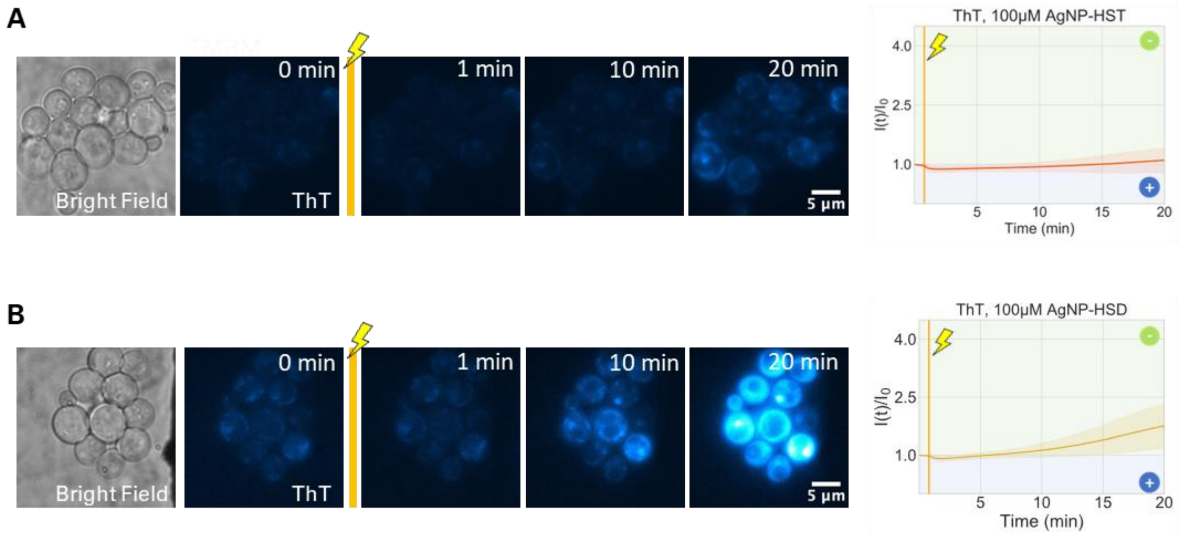


**Fig. S11.** Exposure to biogenic nanoparticles results in responses to EEF that are proportional to the level of the antimicrobial activity the compound exhibits. (A) *S. cerevisiae* cells exposed to AgNP-HST, which possess fungistatic properties. Cells maintain a relative stable membrane potential dynamics in response to EEF. (B) Cells exposed to AgNP-HSD, which does not exhibit significant inhibitory capacity. Cells maintain hyperpolarize in response to EEF.

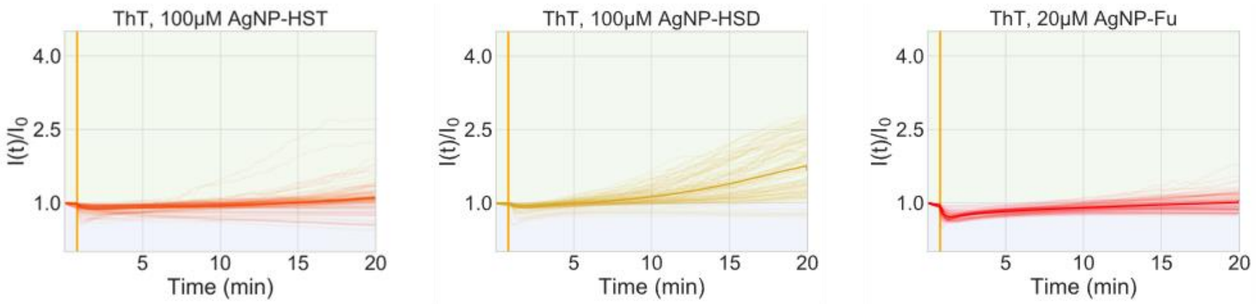


**Fig. S12.** Single-cell time traces of biogenic-nanoparticle-inhibited yeast exposed to EEF, represented by thinner lines. Thicker line represent the mean of single-cell traces.

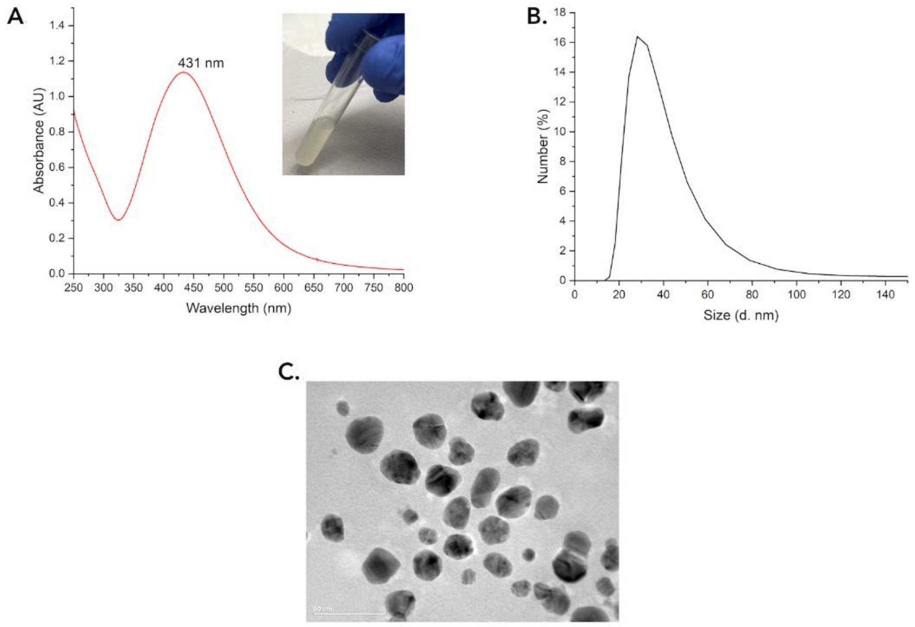


**Fig. S13**. Physical-chemical features of the silver nanoparticles synthesized using hesperidin (AgNP-HSD). A) UV–Vis spectrum of AgNP-HSD in the range of 200-800 nm and the maximum (plasmon band) at 431 nm. Inset is the photographic image of the AgNP-HSD colloid. B) Size distribution and C) Transmission electron microscopy image of AgNP-HSD.

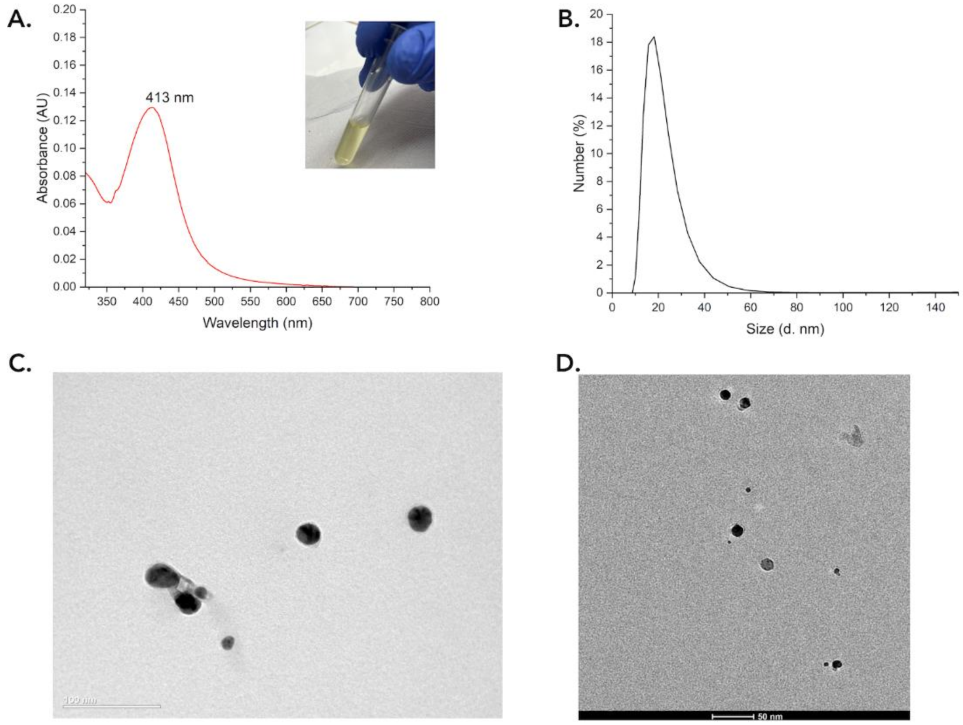


**Fig. S14**. Physical-chemical features of the silver nanoparticles synthesized using hesperetin (AgNP-HST). A) UV–Vis spectrum of AgNP-HST in the range of 325-800 nm and the maximum (plasmon band) at 413 nm. Inset is the photographic image of the AgNP-HST colloid. B) The size distribution, C) Transmission electron microscopy image and D) cryo-EM image of AgNP-HST.

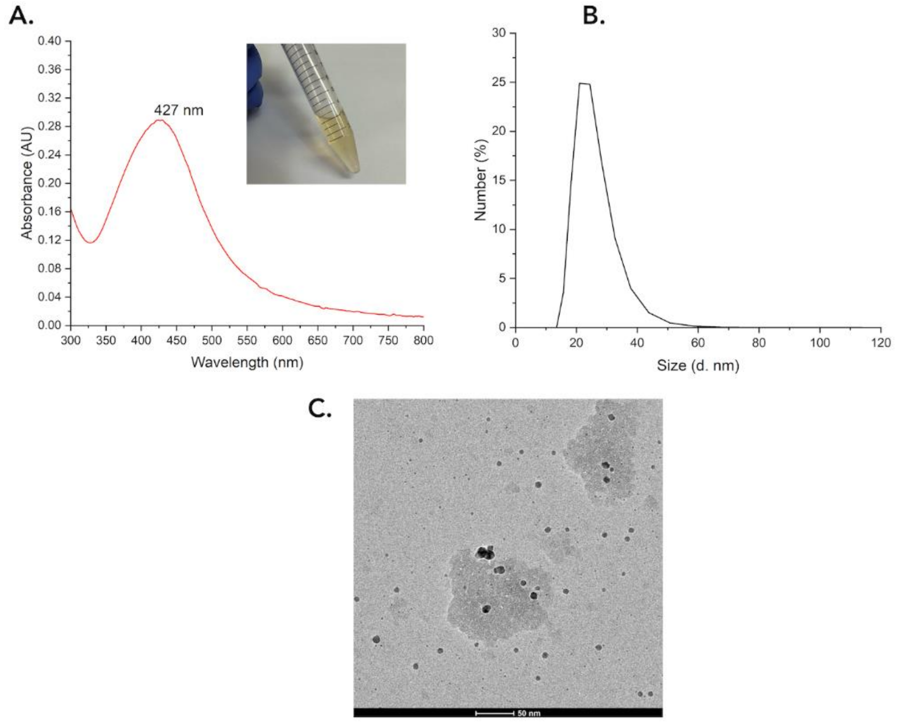


**Fig. S15**. Physical-chemical features of the silver nanoparticles synthesized using fungal filtrate from *Fusarium oxysporum* (AgNP-FU). A) UV–Vis spectrum of AgNP-FU in the range 300-800 nm and the maximum (plasmon band) at 427 nm. Inset is the photographic image of the AgNP-FU colloid. B) The size distribution and C) Cryo-EM image of AgNP-FU.

### Supplementary Table

**Table S1. Characteristics of the synthesized bioAgNPs**

The data obtained from Zeta potential and size measurements (Table S1) indicated that the silver nanoparticles were electrostatically stable and showed medium sizes. The polydispersity indexes indicated that the AgNP colloids were somewhat inhomogeneous regarding the sizes and shapes of the nanoparticles. See also supplementary figures S12 and S13.

| | Average Size (nm) | Zeta Potential (mV) | Polydispersity Index (pdi) |
|---|---|---|---|
| AgNP-HSD | 38.5 | -38.2 + 10.5 | 0.469 |
| AgNP-HST | 22.5 | -38.7 + 10.8 | 0.258 |
| AgNP-FU | 24.7 | -34.5 + 5.4 | 0.476 |

### Supplementary Movies

**Movie S1.** Proliferative cells stained with DiBAC4(3) stimulated by EFF, corresponding to Fig. 2B.
**Movie S2.** Proliferative cells stained with ThT stimulated by EFF, corresponding to Fig. 2C.
**Movie S3.** Proliferative cells stained with TMRM stimulated by EFF, corresponding to Fig. 2D.
**Movie S4.** Yeast cells stained with ThT, exposed to UV, , corresponding to Fig. 3C

### Supplementary Materials and Methods

**Synthesis of AgNP**

Solutions of hesperidin (HSD) or hesperetin (HST), each of 1 mmol $L^{-1}$, were prepared in a sodium hydroxide solution (0.005 mol $L^{-1}$). Then the flavonoid solutions were added dropwise to two flasks with silver nitrate solution of 1 mmol $L^{-1}$ under agitation in a 1:1 (v:v) ratio. After homogenisation, a hydrochloric acid solution of 0.05 mol $L^{-1}$ was added to the nanoparticle colloids until reached pH 7.4. This way, two AgNP colloids were prepared, and named AgNP@HSD and AgNP@HST, respectively.

**UV-Visible spectroscopy**

To initially characterise the nanoparticles an Agilent HP 8453 UV-Visible spectrophotometer was used. The AgNP@HSD and AgNP@HST were placed in quartz cuvettes with a path length of 1 cm. The spectra were taken in the range of 300 to 600 nm. Deionized water was used as a blank.

**Dynamic Light Scattering and Zeta Potential**

To acquire the data a Zetasizer Nano ZS (Malvern Instruments Corp.) analyser was used. The analyses of both Zeta potential and size measurements were made at 25 ºC using a disposable folded capillary cell DTS1070 running three manual measurements with 15 scans each and a balance time of 120 s before beginning.

**Cryogenic Transmission Electron Microscopy**

The diluted samples (1:10, v/v) were put dropped on a sample holder and dried at room temperature. Analyses were performed in a Talos Arctica G2 Cryo-TEM (Thermo Fisher) using HV=120 kV.

**Transmission Electron Microscopy**

AgNP suspensions were diluted in deionized water 1:3 (v/v) and then dried overnight on the sample holder. Analyses were carried out in a Carl-Zeiss Libra 120 microscope using HV=80 kV. Particle counting was done with ImageJ software (National Institute of Health).
